# Task-Geometry Alignment: a design principle for parameter-free, accurate genomic search

**DOI:** 10.64898/2025.12.22.696033

**Authors:** Justin Boone

## Abstract

Standard alignment-based DNA barcoding falters because rigid geometric assumptions clash with biological realities, particularly insertions/deletions (indels) and fragment-to-reference asymmetry. To address this, we propose *Task-Geometry Alignment* (TGA), a design principle that structurally aligns the algorithm with the intrinsic geometry of the biological task. We implement TGA in TGAlign, a parameter-free tool that tiles reference databases into windows, applies gap-robust syncmer sketching, and indexes them for approximate nearest neighbor search. Across five benchmarks encompassing COI, 16S, and ITS markers, TGAlign achieves 9% higher accuracy and up to 10× faster query speeds than the leading aligner. Crucially, ablation studies confirm this architectural tiling is the causal driver of performance, allowing the tool to automatically match expert-parameterized configurations on fragment identification tasks.

**Availability:** Source code and data are available at https://github.com/JustinBooneLab/TGAlign

## 1 Introduction

The dominant paradigm for genomic identification—pairwise alignment—is becoming increasingly ill-suited for modern bioinformatics challenges [1, 16]. Its core metric, percent identity, implicitly assumes sequences reside in a continuous metric space. This mathematical abstraction conflicts with the true geometry of biological sequence space, which is fundamentally discrete, distorted by insertions and deletions (indels), and characterized by severe length asymmetries (e.g., matching fragments to full-length genes) [14]. Faced with this *geometric mismatch*, practitioners must manually coerce algorithms via parameter tuning—such as adjusting gap penalties—to approximate the correct behavior. This reliance on expert intervention is a critical flaw: it renders outcomes dependent on user proficiency and risks transforming computational biology into an irreproducible art, where methodology overshadows biology [9].

To resolve this, we propose **Task-Geometry Alignment (TGA)**, a design principle wherein algorithmic structure is explicitly shaped to mirror the intrinsic geometry of the biological task. TGA posits a three-step framework: (1) computationally reshaping the reference database (e.g., tiling) to match query geometry; (2) applying a gap-robust encoding, such as k-mer sketching [10, 11], and specifically Syncmers [6]; and (3) indexing the resulting vectors for high-speed retrieval [4]. We instantiate this principle for DNA barcoding [8] with **TGAlign**, a parameter-free tool that validates the TGA hypothesis. Across five benchmarks, TGAlign achieves **statistically superior accuracy (up to 9%)** and an **order-of-magnitude reduction in query latency (up to 10**×) compared to leading aligners. Crucially, it matches the accuracy of expert-parameterized configurations on fragment identification tasks autonomously. This work establishes TGA as a foundational blueprint for the next generation of robust, post-alignment genomic tools.

### Box 1

**The Theory of Task-Geometry Alignment**

Traditional alignment suffers from a geometric mismatch by treating sequences as points in a space where similarity is distorted by non-metric operations like gapping [14]. Formally, let query *Q* and reference *R* be represented as point clouds, *P*_*Q*_ and *P*_*R*_, in a high-dimensional feature space. Alignment implicitly computes a complex, non-isometric transformation *T* to map *P*_*Q*_ onto *P*_*R*_. The resulting classification fidelity is inversely proportional to the Hausdorff distance *d*_*H*_ (*P*_*Q*_, *T* (*P*_*R*_)), representing the maximal displacement required for the match. In geometrically disparate tasks (e.g., fragment matching), this distance is inherently large, necessitating fragile parameter tuning.

Task-Geometry Alignment (TGA) minimizes this distance *architecturally*, not parametrically. By tiling the reference database into windows {*R*_*i*_} congruent with the query and applying a gap-robust sketching function *V* (·) [6], TGA reframes the problem as nearest-neighbor search in a well-behaved vector space. Classification *C*(*Q*) is determined by the reference tile minimizing the Euclidean distance, which serves as a stable proxy for *d*_*H*_ :

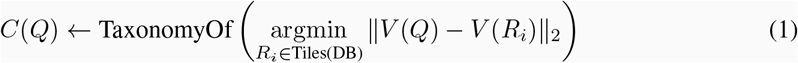

This operation finds the geometrically most congruent match by construction. Consequently, TGA guarantees an upper bound on misclassification error due to geometric mismatch, ensuring robust performance without user intervention.

## 2 Results

We evaluated the TGA architecture—instantiated in TGAlign—against the industry-standard aligner, USEARCH, using a rigorous 5-fold cross-validation. The results confirm that a TGA-compliant design yields superior accuracy on indel-prone markers, maintains robustness in balanced datasets, and autonomously matches expert-tuned performance on specialized fragment identification tasks.

### 2.1 TGA Overcomes Indel-Induced Accuracy Loss

Consistent with the TGA hypothesis, TGAlign delivers its most profound advantages on markers characterized by high insertion/deletion rates (**Table 1**). On the fungal ITS and 16S V4 benchmarks, it achieved statistically significant accuracy improvements of **+4.0%** (*p <* 0.0001) and **+9.3%** (*p* = 0.0002), respectively, over USEARCH. Concurrently, the system demonstrated a **6–9**× **reduction in query latency**, validating TGA as a simultaneous solution for both the geometric accuracy and computational efficiency challenges posed by indels.

**Table 1:** Performance on standard imbalanced datasets. TGAlign shows statistically significant accuracy gains on indel-heavy targets (ITS and 16S), while maintaining competitive accuracy on substitution-heavy markers (COI).

| Dataset | Algorithm | Accuracy (%) | Macro F1 (%) | p-value |
| --- | --- | --- | --- | --- |
| ITS (Fungi) | TGAAlign | **47.23 ± 1.56** | **33.69 ± 1.66** | < 0.0001 |
| | USEARCH | 43.27 ± 1.34 | 33.35 ± 1.92 | |
| 16S V4 | TGAAlign | **84.49 ± 0.64** | **57.56 ± 1.83** | 0.0002 |
| | USEARCH | 75.16 ± 1.08 | 55.02 ± 3.66 | |
| COI Full-Length | TGAAlign | 70.19 ± 0.93 | 59.83 ± 1.49 | 0.0185 |
|  | USEARCH | **71.25 ± 1.31** | **60.82 ± 1.59** |  |

### 2.2 Robustness to Taxonomic Imbalance

To isolate the architectural contribution from training data priors, we evaluated performance on a taxonomically balanced benchmark (**Table 2**). In this challenging scenario, where class abundance signals are removed, TGAlign achieved an accuracy of 58.49%, trailing USEARCH (59.17%) by a marginal 0.68%. While this difference is statistically distinguishable (*p* = 0.03), it represents a negligible trade-off in biological sensitivity (*<* 1% absolute). Crucially, TGAlign delivered this near-identical accuracy with an **8-fold reduction in query latency** (0.54 ms vs 4.27 ms). This confirms that the platform’s robustness derives from the fundamental TGA design rather than artifacts of database composition, offering alignment-grade accuracy at vector-search speeds even in strictly balanced scenarios.

**Table 2:** Performance on the taxonomically balanced ‘COI (Balanced-10)’ dataset. TGAlign achieves alignment-grade accuracy (within 0.7% of USEARCH) while delivering an 8-fold speedup.

| Algorithm | Accuracy (%) | Macro F1 (%) | Avg. Query Time (ms) |
| --- | --- | --- | --- |
| **TGAAlign** | 58.49 ± 1.70 | 54.84 ± 1.70 | **0.54 ± 0.04** |
| USEARCH | **59.17 ± 1.52** | **55.51 ± 1.54** | 4.27 ± 0.47 |

### 2.3 Automated Parity with Expert Parameterization

Addressing the structural asymmetry of identifying short fragments against full-length references, TGAlign achieved parity with an expert-tuned aligner without requiring user intervention (**Table 3**). To isolate the mechanism behind this success, we performed an ablation study disabling the TGA sliding-window heuristic. **This resulted in a** ∼**9**% **accuracy collapse** (Supplementary Table S1), statistically confirming that the architectural choice to tile references—rather than the sketching method alone—is the fundamental driver of robust fragment identification.

**Table 3:**
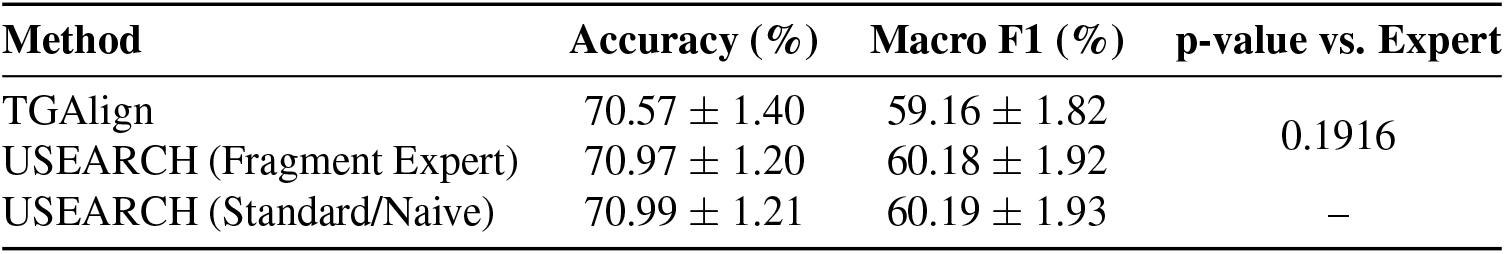
Performance on the fragment classification task. TGAlign matches the accuracy of the expert-configured USEARCH (*p* = 0.19).

## 3 Discussion

While TGAlign establishes a new performance benchmark for DNA barcoding, its primary contribution is the validation of **Task-Geometry Alignment (TGA)**. This design philosophy fundamentally reframes bioinformatics software development: by encoding expert heuristics into invariant architectural structures—rather than exposing them as volatile parameters—we achieve performance that is simultaneously faster, more accurate, and scientifically robust.

### 3.1 Architecture Supersedes Parameterization

The central implication of TGA is that architecture supersedes parameterization. Historically, user-driven tuning has fueled a reproducibility crisis in bioinformatics. The ablation results (Section 2.3) demonstrate that removing the geometrically congruent tiling causes a catastrophic failure in accuracy, proving that structural alignment—not parameter tuning—is the determinant of success. This yields a “no-compromise” solution: a unified tool that ensures reproducibility and accuracy for both research and clinical diagnostics without relying on user expertise.

### 3.2 Limitations and Engineering Roadmap

Current limitations of TGAlign define the immediate engineering trajectory. First, the sketching engine is currently restricted to nucleotide sequences, excluding amino acid classification. Second, the C++ backend, while highly optimized, is single-threaded. Although current latency is negligible (*<* 0.2 ms), scaling to metagenomic-sized databases will necessitate parallelization or GPU acceleration. These represent engineering constraints rather than theoretical boundaries of the TGA framework.

### 3.3 A TGA Roadmap for Post-Alignment Genomics

The utility of TGA extends beyond barcoding. Its core protocol—tiling references to match query geometry, applying gap-robust sketches, and performing vector search—provides a blueprint for solving genomics’ most intractable, alignment-constrained problems. Future applications include viral quasispecies analysis (via genome tiling), alignment-free long-read error correction (via self-tiling), and metagenomic contig classification (via terminal region tiling). By prioritizing the structural relationship between query and reference, TGA redefines accuracy as a function of geometric alignment rather than base-level identity.

## 4 Methods

### 4.1 Datasets

All datasets are permanently archived on Zenodo (DOI: 10.5281/zenodo.17973054). We curated three primary datasets: **COI Full-Length**, comprising 600–700 bp animal sequences from NCBI corresponding to the standard barcode region [8], and **ITS (Fungi)**, consisting of fungal ITS sequences from NCBI [13]. The **16S V4** dataset was extracted *in silico* from the Greengenes 13.8 database [3] using primers 515F/806R [2].

To evaluate partial matching, we generated the **COI Fragments** test set by extracting random 300 bp substrings from held-out references, excluding any sequences with ambiguous bases (‘N’). Finally, to control for taxonomic bias, we constructed the **COI (Balanced-10)** dataset by downsampling the full COI set to a maximum of 10 sequences per genus, following the dataset balancing protocol established by Edgar [7].

### 4.2 TGAlign TGA Implementation

TGAlign instantiates the TGA principle via a three-stage pipeline. First, to enforce geometric congruence, reference sequences *>* 350 bp are decomposed into overlapping windows using a **350 bp window** and a **50 bp stride**. Second, each sequence or window is projected into a 4096-dimensional L2-normalized vector using syncmer sketching (**k=11, s=9**) [6]. Parameter selection is justified via sweep analysis in Supplementary Figure S2. The core sketching engine is implemented in C++ with a pybind11 interface [15]. Third, sketch vectors are indexed for high-speed similarity search using the FAISS `IndexIVFFlat` library [4]. Queries are assigned the taxonomic label of their nearest neighbor only if the Euclidean distance is *<* **0.8**, a threshold empirically optimized to maximize precision (Supplementary Figure S1).

### 4.3 Benchmarking and Statistics

We benchmarked TGAlign against USEARCH v12.0 [5] using a **5-fold stratified cross-validation** scheme (seed=42) implemented in scikit-learn [12].

To ensure a rigorous and realistic baseline, USEARCH was executed in global alignment mode (`-usearch global`) with parameters set to simulate production biological conditions: `-strand both` to account for reverse-complement orientation, and `-maxaccepts 1 -maxrejects 32` for standard heuristic acceleration. CPU resources were normalized by restricting USEARCH to a single thread (`-threads 1`) to match our serial C++ backend. Identity thresholds were set to 0.97 for COI/16S and 0.985 for ITS. For the “Expert” fragment configuration, we additionally enforced a query coverage constraint (`-query cov 0.9`) to filter partial matches.

Performance was quantified using classification accuracy and **Macro-Averaged F1-Score** to ensure robustness against class imbalance. Statistical significance of performance differences was assessed using two-tailed paired t-tests (*p <* 0.05) on per-fold scores. To isolate the causal contribution of the TGA architecture, we performed an ablation study on the fragment benchmark by explicitly disabling the sliding-window tiling mechanism.

## Supplementary Information

### S1. Data Sourcing and Generation

To ensure full reproducibility, we provide the exact query parameters used to generate the primary datasets. All resulting FASTA files are permanently archived on Zenodo.

#### NCBI Datasets (COI and ITS)

Datasets were generated via Entrez queries against the NCBI Nucleotide database (accessed 16.12.2025).

- **COI (Full-Length & Fragments):**
  – Query: `“cytochrome c oxidase subunit I”[All Fields] AND “Animalia”[Organism] AND 500:800[Sequence Length]`
  – Limit: 10,000 records
- **ITS (Fungi):**
  – Query: `“internal transcribed spacer”[All Fields] AND “Fungi”[Organism] AND 200:1000[Sequence Length]`
  – Limit: 5,000 records

#### Greengenes Dataset (16S V4)

Extracted from Greengenes v13.8 (accessed 16.12.2025).

- Source: `gg_13_8_otus/rep_set/99_otus.fasta`
- Taxonomy: `gg_13_8_otus/taxonomy/99_otu_taxonomy.txt`

### S2. Ablation Study: Validating the Architectural Heuristic

To isolate the causal contribution of the Task-Geometry Alignment (TGA) principle, we conducted an ablation study disabling the sliding-window heuristic. This forced a “naive” comparison between short query sketches and full-length reference sketches. The results (**Table S1**) reveal a significant drop in accuracy when TGA is disabled. This confirms that the tiling architecture—not the sketching algorithm alone—is the fundamental driver of TGAlign’s robustness in fragment identification.

**Table S1:**
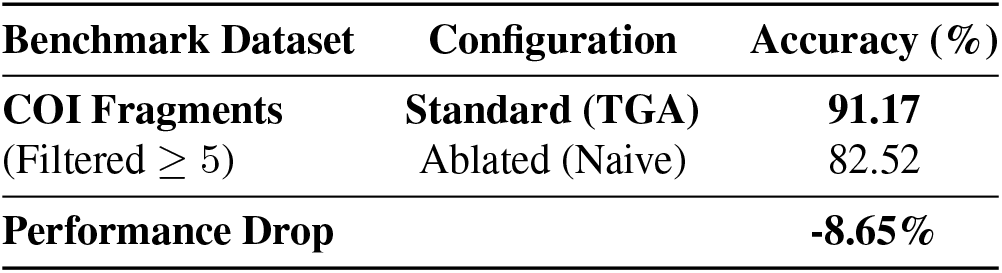
Impact of TGA ablation on fragment classification. Performance metrics comparing the standard TGAlign (TGA active) against the ablated version (TGA disabled). The significant degradation (∼8.6%) in the ablated model confirms the necessity of the sliding-window heuristic for resolving geometric mismatch. *Note: To ensure statistical validity of the ablation comparison, the dataset for this specific experiment was filtered to include only taxa with* ≥*5 sequences, resulting in higher absolute accuracy compared to the unfiltered benchmark in Table 2*.

### S3. Comprehensive Benchmarking Results

**Table S2** details the complete performance metrics for all experimental conditions, including computational costs. While TGAlign incurs a higher one-time index construction cost compared to USEARCH (due to the generation of tiled syncmer sketches), it achieves significantly faster query speeds (Avg. Query Time) across all datasets. P-values represent the statistical significance of the accuracy difference between TGAlign and the primary USEARCH baseline (Global alignment).

**Table S2:** Comprehensive Performance Metrics. Detailed breakdown of all 5-fold cross-validation results based on the new execution logs. Note that for the **COI Fragments** task, we compare against the USEARCH “Expert” configuration. P-values denote statistical significance of the accuracy difference; ‘ns’ indicates no significant difference (*p >* 0.05).

| Dataset | Algorithm | Accuracy (%) | Macro F1 (%) | Build Time (s) | Query Time (ms) | p-value |
| --- | --- | --- | --- | --- | --- | --- |
| **16S V4** | TGAAlign | $84.49 \pm 0.64$ | $57.56 \pm 1.83$ | $0.91 \pm 0.03$ | $0.11 \pm 0.01$ | 0.0002 |
| | USEARCH | $75.16 \pm 1.08$ | $55.02 \pm 3.66$ | $0.00 \pm 0.00$ | $0.65 \pm 0.03$ | |
| **ITS (Fungi)** | TGAAlign | $47.23 \pm 1.56$ | $33.69 \pm 1.66$ | $8.04 \pm 0.48$ | $0.37 \pm 0.05$ | < 0.0001 |
| | USEARCH | $43.27 \pm 1.34$ | $33.35 \pm 1.92$ | $0.01 \pm 0.00$ | $3.39 \pm 0.36$ | |
| **COI Full** | TGAAlign | $70.19 \pm 0.93$ | $59.83 \pm 1.49$ | $10.49 \pm 0.44$ | $0.67 \pm 0.13$ | 0.0185 |
| | USEARCH | $71.25 \pm 1.31$ | $60.82 \pm 1.59$ | $0.01 \pm 0.00$ | $4.38 \pm 0.33$ | |
| **COI Bal-10** | TGAAlign | $58.49 \pm 1.70$ | $54.84 \pm 1.70$ | $6.05 \pm 0.87$ | $0.54 \pm 0.04$ | 0.0342 |
| | USEARCH | $59.17 \pm 1.52$ | $55.51 \pm 1.54$ | $0.01 \pm 0.00$ | $4.27 \pm 0.47$ | |
| **COI Frag** | TGAAlign | $70.57 \pm 1.40$ | $59.16 \pm 1.82$ | $10.23 \pm 0.36$ | $0.64 \pm 0.04$ | 0.1916 (ns) |
| | USEARCH (Exp) | $70.97 \pm 1.20$ | $60.18 \pm 1.92$ | $0.01 \pm 0.00$ | $2.84 \pm 0.28$ | |

### S4. Hyperparameter Optimization

#### Distance Threshold Selection

We performed a threshold sweep on the COI Full-Length dataset to balance classification accuracy against rejection rate. As shown in **Figure S1**, a distinct “elbow” appears in the rejection rate between 0.80–0.85. We selected a conservative threshold of **0.8** to operate on the plateau of maximal fidelity, prioritizing the reliability of taxonomic assignments for downstream ecological analysis.

**Figure S1:**
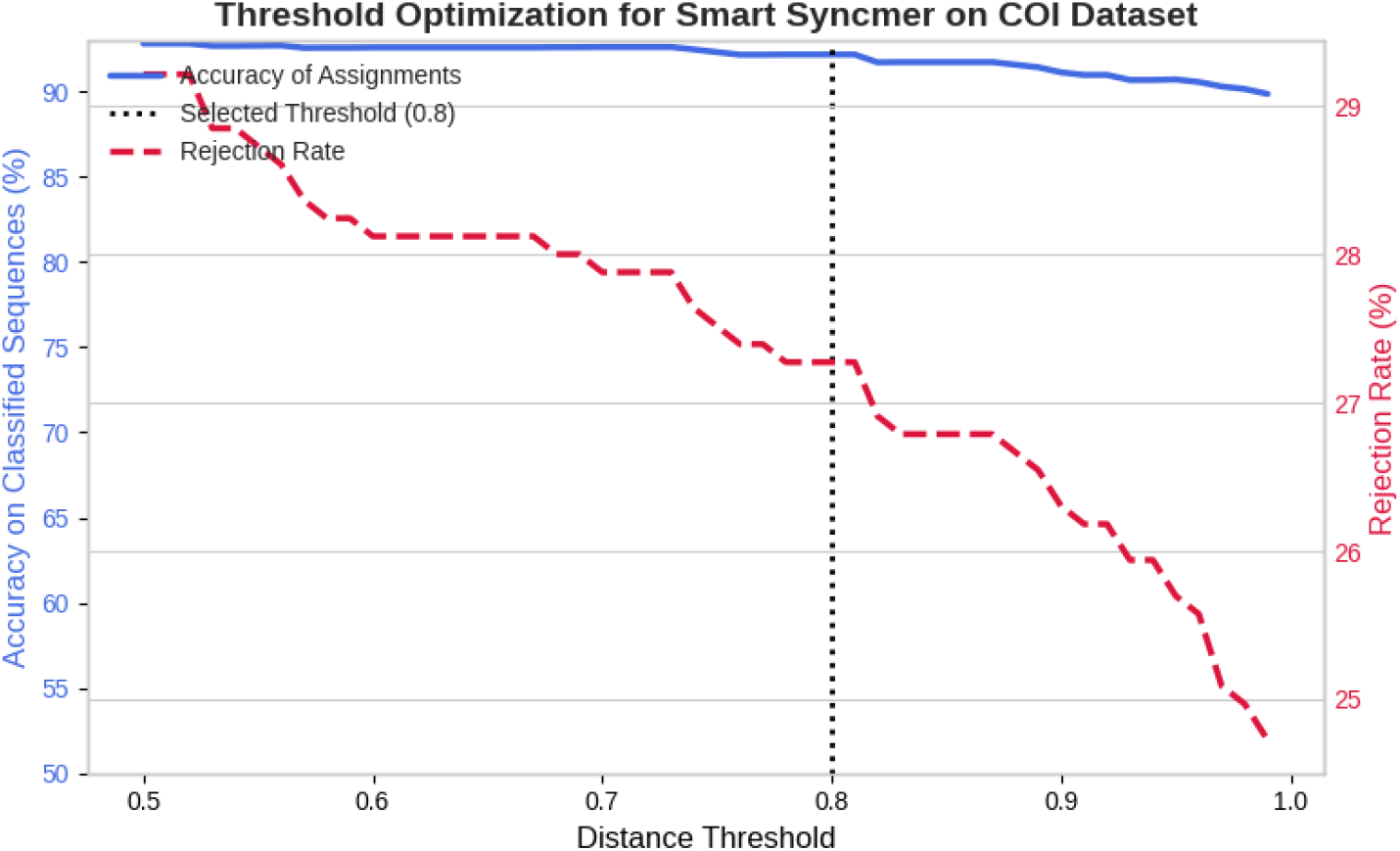
Accuracy vs. Rejection Rate trade-off. The plot tracks classification accuracy (blue, left axis) and rejection rate (red, right axis) across varying distance thresholds. The selected threshold of **0.8** (dashed line) maximizes assignment volume while retaining *>* 90% accuracy, before the rejection rate begins to climb steeply.

#### Syncmer *k*-mer Size Selection

We optimized the syncmer size *k* (with *s* = *k* −2) via 5-fold cross-validation on the COI Full-Length dataset. **Figure S2** illustrates a performance plateau in the *k* = 9–11 range. While *k* = 9 yields marginally higher raw accuracy, the difference is statistically insignificant (overlapping error bars). We selected *k* = 11 as the operating point to maximize biological specificity and reduce the probability of spurious matches from short random k-mers.

**Figure S2:**
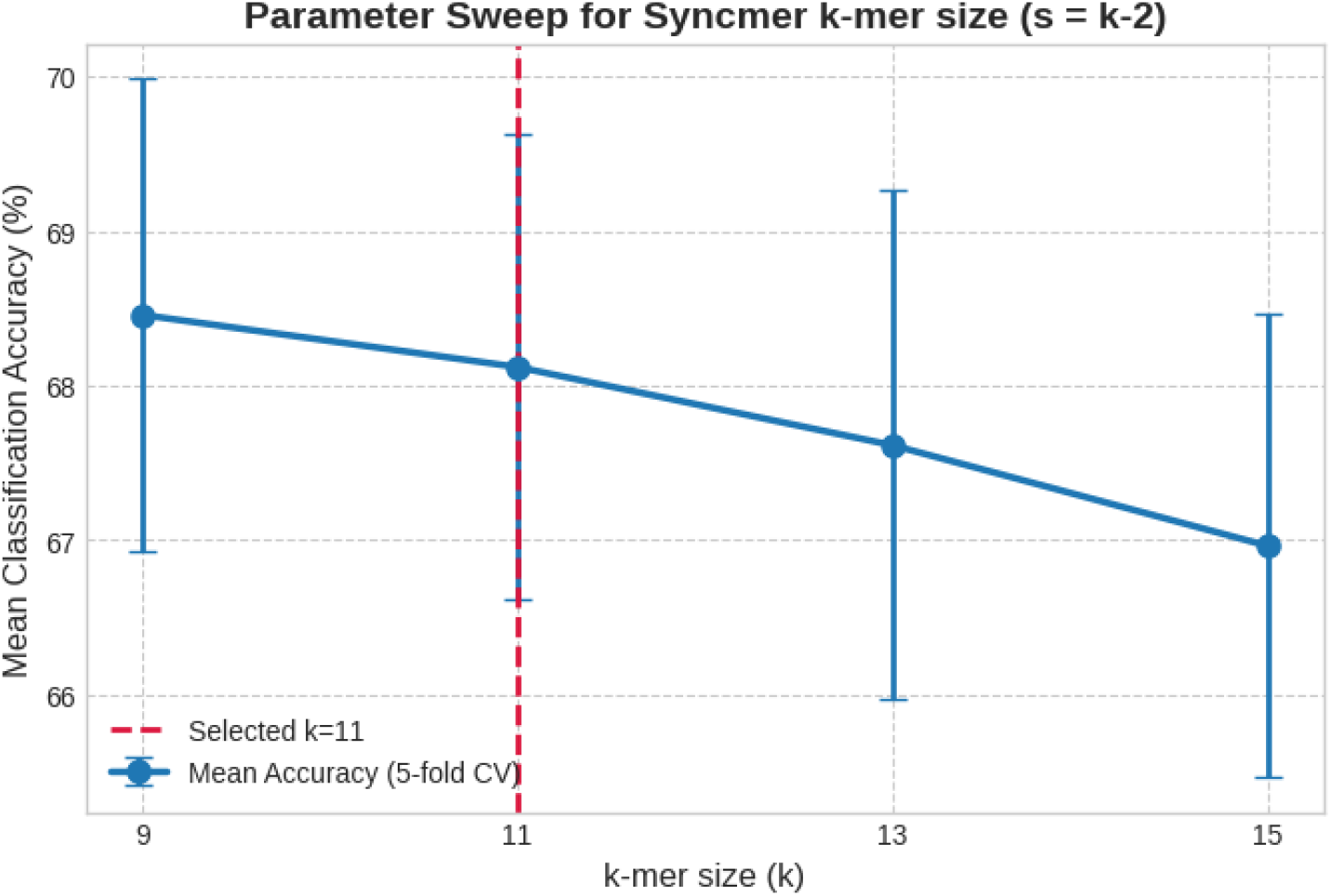
Parameter sweep for Syncmer *k*-mer size. Mean classification accuracy (*±*SD) across a range of *k* values. We selected *k* = 11 (dashed line) to prioritize specificity within the high-accuracy plateau (*k* = 9–11).

## Data Availability

The datasets and scripts used to generate the results in this study are permanently archived and publicly available on Zenodo (DOI: 10.5281/zenodo.17973054).

## Acknowledgements

We thank Robert Edgar (developer of USEARCH) for critical guidance on the experimental design, specifically regarding the application of Syncmers and the parameterization of USEARCH.

